# Network analysis of mass spectrometry imaging data from colorectal cancer identifies key metabolites common to metastatic development

**DOI:** 10.1101/230052

**Authors:** Paolo Inglese, Nicole Strittmatter, Luisa Doria, Anna Mroz, Abigail Speller, Liam Poynter, Andreas Dannhorn, Hiromi Kudo, Reza Mirnezami, Robert D Goldin, Jeremy K Nicholson, Zoltan Takats, Robert C Glen

## Abstract

A deeper understanding of inter-tumor and intra-tumor heterogeneity is a critical factor for the advancement of next generation strategies against cancer. The heterogeneous morphology exhibited by solid tumors is mirrored by their metabolic heterogeneity. Defining the basic biological mechanisms that underlie tumor cell variability will be fundamental to the development of personalized cancer treatments. Variability in the molecular signatures found in local regions of cancer tissues can be captured through an untargeted analysis of their metabolic constituents. Here we demonstrate that DESI mass spectrometry imaging (MSI) combined with network analysis can provide detailed insight into the metabolic heterogeneity of colorectal cancer (CRC). We show that network modules capture signatures which differentiate tumor metabolism in the core and in the surrounding region. Moreover, module preservation analysis of network modules between patients with and without metastatic recurrence explains the inter-subject metabolic differences associated with diverse clinical outcomes such as metastatic recurrence.

**Significance:** Network analysis of DESI-MSI data from CRC human tissue reveals clinically relevant co-expression ion patterns associated with metastatic susceptibility. This delineates a more complex picture of tumor heterogeneity than conventional hard segmentation algorithms. Using tissue sections from central regions and at a distance from the tumor center, ion co-expression patterns reveal common features among patients who developed metastases (up of > 5 years) not preserved in patients who did not develop metastases. This offers insight into the nature of the complex molecular interactions associated with cancer recurrence. Presently, predicting CRC relapse is challenging, and histopathologically like-for-like cancers frequently manifest widely varying metastatic tendencies. Thus, the methodology introduced here more robustly defines the risk of metastases based on tumor biochemical heterogeneity.

**Author contributions:** P.I., Z.T., R.C.G.: designed the study, developed the workflow, analyzed the data, interpreted the results, wrote the paper; N.S. collected the MS, performed the H…E staining, wrote the paper; L.D.: interpreted the results, wrote the paper; A.M.: collected the MS; A.S.: histological assessment; L.P.: collected the tissue specimens and clinical metadata; A.D.: collected the MS; H.K.: performed the H…E staining; R.M.: collected the tissue specimens and clinical metadata. R.G.: histological assessment; J.K.N: designed the study, edited the paper.

## Introduction

Cancerous tissue is characterized by a high level of heterogeneity when compared to the surrounding host tissue. This heterogeneity, expressed at multiple levels, may be purely genetic (Burrell, McGranahan et al. 2013) or epigenetic (Marusyk, Almendro et al. 2012) or a combination, and represents one of the most challenging aspects for the implementation of effective anti-cancer treatment strategies. In earlier work, mass spectrometry imaging (MSI) techniques have shown promise in their ability to capture the diversity of tumors through the analysis of molecular expression patterns (Schwartz, Weil et al. 2004, Eberlin, Dill et al. 2010, McDonnell, Corthals et al. 2010, Schwamborn and Caprioli 2010, Eberlin, Norton et al. 2012, Alexandrov, Becker et al. 2013), showing that the underlying heterogeneity is reflected in dramatic metabolic changes across different tumor sub-regions. From the point of view of statistical modelling, the lack of detailed metabolic data of the heterogeneous properties of cancerous tissues and the difficulty of detecting those differences through visual inspection of the specimens makes necessary the use of analytical methods combined with unsupervised analysis techniques. These approaches can detect the underlying statistical structure of the data and can be employed to cluster molecular abundance patterns that are found to be common among discrete regions of tissues. These subsets can be defined both in the spatial domain (pixels) and in the metabolic domain (ions corresponding to molecules) and both approaches have been explored in previous work for the study of cancer and other types of tissues. Linear methods, such as PCA or MDS, combined with hierarchical clustering have shown that MALDI-MSI can capture the heterogeneity of cancer while allowing analysis of the properties of the metabolic space (Deininger, Ebert et al. 2008, Rauser, Marquardt et al. 2010), whereas other approaches based on the combination of multiple clustering methods base their results on the presence of concordance between the spatial patterns associated with clusters (pixels) (Jones, van Remoortere et al. 2011, Balluff, Frese et al. 2015). Other studies have associated the presence of specific sets of molecules in the cancerous tissue with clinical outcome (e.g. increased risk of mortality or metastasis development) (Abdelmoula, Balluff et al. 2016, Lou, Balluff et al. 2016). However, the conclusions from this earlier work are often limited to providing a series of possible biomarkers for tumor heterogeneity - limited or no insight is given into the possible biochemical mechanisms related to the increased/decreased abundance of identified markers in a specific region of the tissue.

We hypothesize that the mechanisms behind tumor heterogeneity are much more complex than those captured simply by the identification of sets of locally highly abundant metabolites and, for this reason, it is extremely important to introduce methodologies capable of capturing a broader view of the underlying biochemical interactions. Network analysis represents a natural choice in this regard, since it provides a set of tools to describe the possible interactions (co-expressions) between variables in terms of their mutual statistical similarities. In the present study, we have characterized the metabolic differences between patients with or without metastatic recurrence using weighted coexpression network analysis (WGCNA) (Langfelder and Horvath 2008) module preservation (Langfelder, Luo et al. 2011). Previous studies have applied a WGCNA approach to metabolic data (DiLeo, Strahan et al. 2011, Su, Wang et al. 2014, Yu, Niu et al. 2015) and, specifically, several studies have made use of module preservation as a measure for the identification of (gene) expression differences between species or disease conditions (Oldham, Horvath et al. 2006, Miller, Horvath et al. 2010, Horvath, Zhang et al. 2012, Chen, Cheng et al. 2013, Tong, Li et al. 2013, Xue, Huang et al. 2013, Huang, Maruyama et al. 2014) but, to the best of our knowledge, this is the first application to MSI data derived from cancerous tissue specimens. A related study uses correlations to evaluate the ion co-localization in MSI data (McDonnell, van Remoortere et al. 2008). The hypothesis underlying the present study is that network modules can represent different aspects of the local metabolism through the analysis of co-localized ions and that the metabolic differences characterizing tumor aggressiveness can be identified by those modules associated with patients with metastases that are (or are not) preserved in patients without metastases.

For both the data collected at the tumor core and at a distance of 10cm from the tumor, two groups of patients are defined: a *control cohort* consisting of patients with metastatic recurrence and a *test cohort* consisting of those patients without metastatic recurrence. Using WGCNA, an ion network can be constructed using the mass spectral data from instances where the variables have stronger connections if co-localized (correlated). However, in contrast to gene expression analysis, where each sample can be treated individually, MSI data from a single patient specimen often consists of thousands of mass spectral profiles (one for each pixel). This, together with the different number of patients belonging to the two cohorts, may result in a significantly different number of spectra associated with the two clinical outcomes. Furthermore, treating the spectra as independent measures does not guarantee that the correlations reflect the co-expression levels of the ions within each individual tissue specimen.

For these reasons, a comparative analysis is performed using a consensus cohort network, defined by merging the tissue sample networks from a single cohort. A set of modules are defined by partitioning the consensus network into sub-networks of highly correlated nodes and the metabolic differences between the two cohorts are characterized in terms of module preservation.

In this study, the WGCNA module preservation analysis on DESI-MSI data from specimens surgically removed at the tumor center and at a distance of 10cm from the tumor center (Table 1) shows, using no prior information about the nature of the local tissue (tumor, surrounding healthy tissue or background), that the metastasis associated metabolic patterns within the tumor core microenvironment involve not only the cancerous tissue, but are extended to the neighboring stromal tissue and furthermore, that metastasis associated differences (albeit different ones) in the cellular metabolism are also detected in regions relatively distant from the tumor.

**Table 1.**
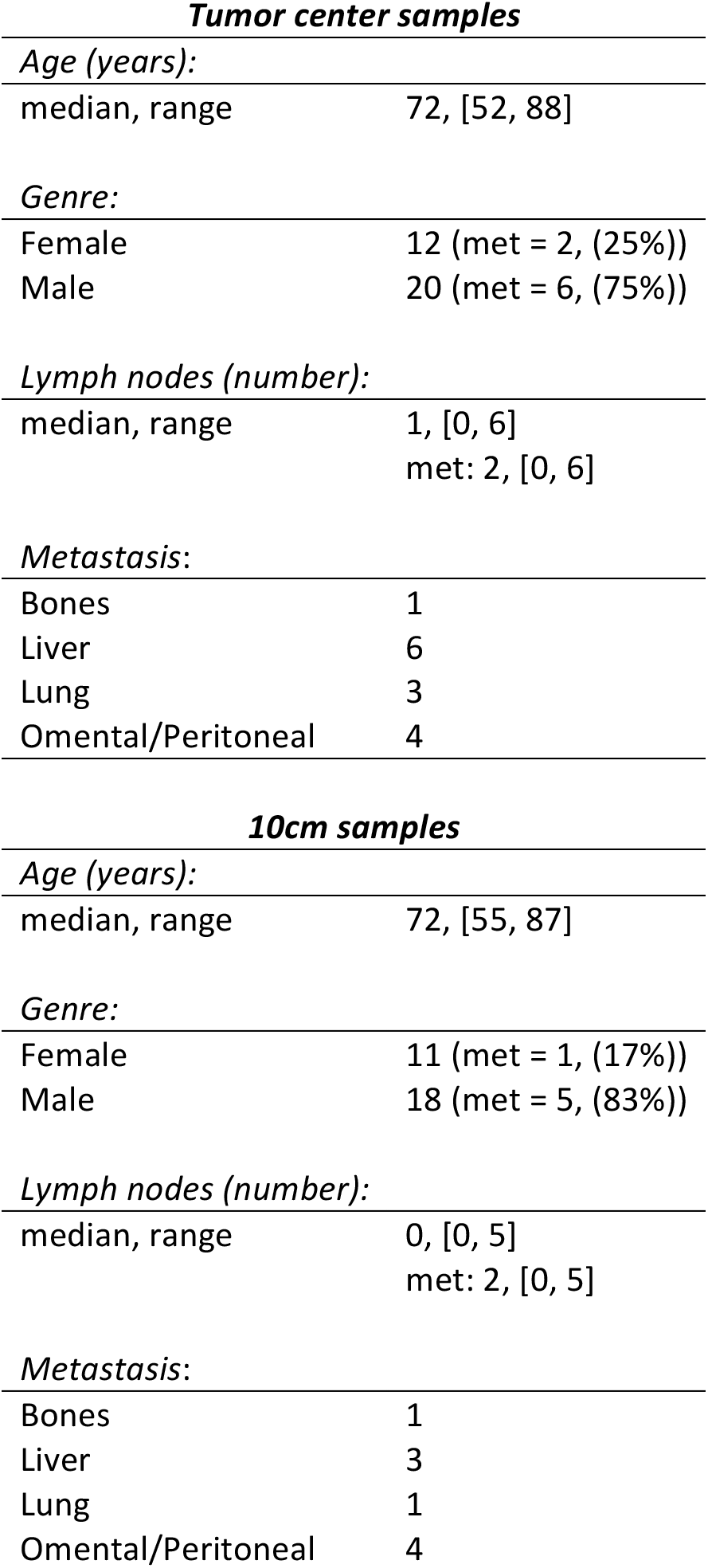
– Patient clinical metadata associated with the two datasets.

## Results

### Re-calibration of peak positions, visual assessment of sample quality and ion selection

The peak lists (centrode data), extracted from the RAW data using the ProteoWizard software (Kessner, Chambers et al. 2008) were re-calibrated in order to reduce the peak shifts arising from specific instrumental conditions during the MS acquisition. Since a dedicated reference compound for lock-mass correction was not infused during the MS measurement process, a re-calibration procedure was introduced exploiting the characteristics of the analyzed data. Two ions were identified as reference candidates because of their standard abundance across all the samples: palmitate ion, corresponding to 255.2330 *m/z* ([M-H]^−^), was found homogeneously occurring in the entire layer containing both the tissue and the background; whereas phosphatidylinositol, PI(38:4), corresponding to 885.5499 *m/z* ([M-H]^−^) was mainly found in the region occupied by the tissue. Using these two reference ions, the *m/z* shift across the entire sample could be estimated. Setting a 1 *Δppm* wide search window, the intensity of the closest peak to the reference ions was associated with each pixel; if no peak was found within the search window, the pixel was left blank. Because of the characteristics of the two reference ions, it was expected that the intensity of the image corresponding to palmitic acid would cover the entire layer whereas the image corresponding to PI(38:4) would clearly represent the tissue sample. If the value of *Δppm* was too small, the matched ions intensity image would instead present holes or scattered pixels. In this way, after scanning increasing values of Δ *ppm* in steps of 1ppm, the optimal *Δppm* was defined as the minimum value to produce images defined as having no more than 1% of missing pixels for the palmitic acid image and a clear tissue image for PI(38:4) (Fig. 1A-F). If the optimal *Δppm* was larger than 10ppm (more than double the instrumental error), then the quality of the sample was considered insufficient and the sample was discarded. Subsequently, the *m/z* values found in the search window corresponding to the palmitate peak were used to quantify the peak shift across all the pixels. A robust Locally Weighted Scatterplot Smoothing (LOWESS) model was used to estimate the relative distance between the RAW peak positions and the expected *m/z* value (255.2330 *m/z)* (Fig. 1G). Only the palmitic acid was used to estimate the peak shifts; for this ion, the LOWESS model was also capable of estimating the peak shift in the presence of a small number of blank pixels, whereas it failed to estimate the peak shift for PI(38:4) because of the large number of consecutive blank pixels (Fig. 1H). The estimated peak shift was used to re-calibrate all the spectra, applying a rigid translation of the *m/z* values, so that the *m/z* values corresponding to palmitic acid were set equal to the theoretical value. While this approach gave satisfactory results for Orbitrap data, the same algorithm cannot be directly used for the correction of time-of-flight data.

**Fig. 1.**
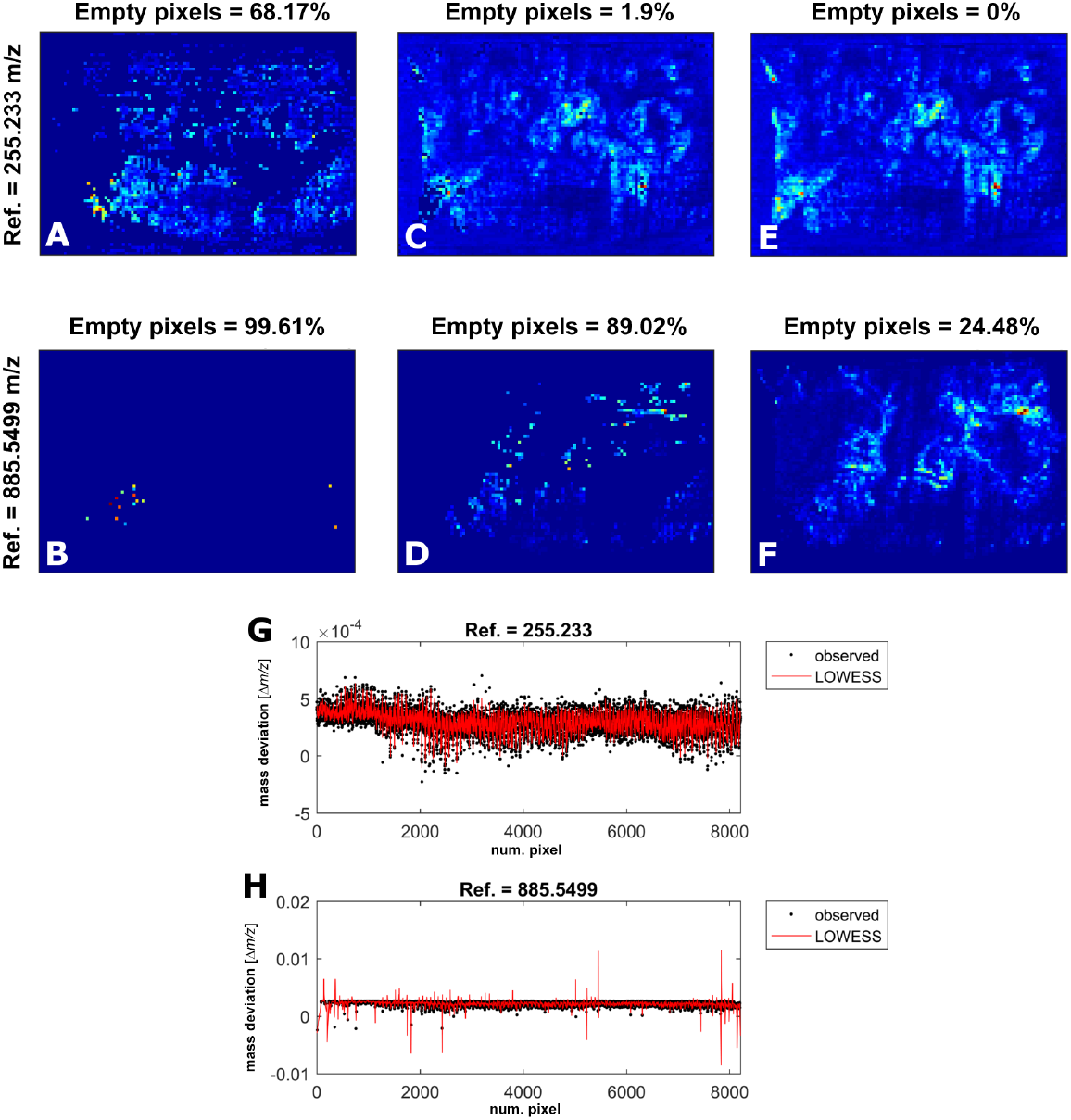
*– Effect of the search wi"dow width (Δppm) o" the peaks detected i" the raw data. In the example shown in the figure, a Δppm equal to 1 or 2 is too small compared to the maximum peak shift found i" the sample, producing scattered images for both the palmitic acid and PI(38:4) (Δppm = 1, A-B; Δppm = 2, C-D). In contrast, using the value Δppm = 3, a peak corresponding to 255.2330 m/z for the full layer (0% empty pixels, E) can be detected and produced a" image similar to the stained tissue sample (24.48% empty pixels, F). The robust LOWESS model fitted to the peak shift measured with Δppm = 3 captures the m/z shift across all the pixels (G), whereas it fails to estimate the m/z shift for the peaks corresponding to 855.5499 m/z because of the large number of missing pixels and also because of the gaps related to the localized presence of tissue i" the image (H)*.

After peak matching (SI Appendix, Materials and Methods), in order to compare the different cooccurrence patterns, only the ions present in all the samples in at least 1% of the pixels were used. Additionally, in order to consider only the biological correlation patterns, isotopes were identified and removed. An ion at an *m m/z* position was denoted as an isotope if there existed another ion at *m*_0_ in the matched ions list such that *m* ∈ [*m*_0_ — 1.003 *k,m_0_ +* 1.0045 *k*], with *k* ∈ ℕ and such that it could not be annotated using an error threshold of 5 ppm were checked.

### Modelling the biochemical differences between groups of patients through the non-preserved network ion modules

In all experimental analyses, two patients’ cohorts were defined: the *reference cohort, X_met_*, consisting of those patients that showed clinical outcomes of interest (developed metastasis), and the *test cohort, X_non-met_*, consisting of those patients who did not develop metastasis during the follow-up period. The assumption behind this procedure was that the differences in ion abundances in local regions of the tissue reflected metabolic heterogeneity and that differences in metabolic pathways occurring in the cancer cells were associated with a different clinical outcome. In order to calculate a representative network for each cohort, the definition of a consensus network was used (Langfelder and Horvath 2008). A signed adjacency (SI Appendix, Materials and Methods) was calculated from each tissue specimen of the cohort, and combined into a single matrix whose elements were defined as the corresponding minimum value across all the cohort samples. The soft power *β*, necessary to calculate the signed topological overlap matrix (TOM) was determined as the smallest integer corresponding to a *R^2^ >* 0.8 *(R^2^ > 0.7* for 10cm samples) for the scale-free network assumption for both *X_met_* and *X_non-met_* in a range of 1 to 20. Using the same value of *β* aimed to preserve the compatibility between the control and test networks. In a similar way, the consensus TOM from the only control data, denoted *TOM_met_*, was defined as the matrix whose elements were equal to the corresponding minimum values across the *TOMs* of the cohort (SI Appendix, Materials and Methods).

In order to obtain a partition of disjoint sets of highly correlated ions (denoted *reference modules)* in the reference network, a hierarchical clustering with average linkage on the *TOM_met_* based distance matrix was applied. The distance matrix, calculated as *1-TOM_met_*, represented the dissimilarities between the ion expressions within the metastatic tissue specimens. The optimal partition was identified using the Dynamic Tree Cut hybrid algorithm (Langfelder, Zhang et al. 2008) with a minimum number of ions per cluster equal to 5. The latter choice was based on the hypothesis that small clusters could capture a more detailed picture of the spatial similarities in the molecular expressions. The module eigenmetabolites (ME) (in a similar fashion to the module eigengenes of the original WGCNA algorithm, these were defined as the scores of the first principal component of the MSI data limited to the module ions) defined the representative spatial distribution of the ions belonging to the module. In order to reduce the potential redundancy of similar ion modules, those modules where the MEs were characterized by a Pearson’s correlation larger than 0.8 were merged into a single module. The ions that were not clustered (because they were assigned to modules with less than 5 ions) were assigned to the “grey” module and were not passed to the next stage of the analysis. In this way, a set of modules were associated to the consensus network, representing the observed molecular heterogeneity of the cancerous tissue. The metabolic differences between the reference cohort of patients and test cohort of patients were determined by module preservation analysis (SI Appendix, Materials and Methods). The *X_met_* consensus network modules were tested for preservation in the *X_non--met_* consensus network, under the hypothesis that not-preserved modules would capture the metabolic differences associated with the patients’ prognosis.

### Phosphatidylglycerol metabolism in stromal tissue surrounding the tumor core is associated with metastasis

The mass spectral data from the tumor core specimens of 32 subjects (Table 1) was used to perform the module preservation analysis. The MSI data from the 8 patients that developed metastasis, denoted 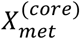, was used to define the consensus reference network, while the remaining 24 patients’ MSI data, denoted 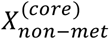, was used to build the test network. The high correlation between their node connectivity *k*, defined as the sum of the node edges strengths (Oldham, Horvath et al. 2006), (Pearson’s r = 0.95, Fig. 2A) showed that the two networks were characterized by similar global properties. This result showed that the global tumor metabolism was undifferentiated between the two groups of patients. By partitioning the 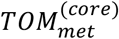 using the hierarchical clustering, 6 reference modules were found (denoted as “blue”, “brown”, “green”, “red”, “turquoise” and “yellow”) (Fig. 2A). A visual inspection of the ME images showed that 4 modules were associated with the tissue and 2 were localized mainly outside of the tissue (Fig. S2). Moreover, it was seen that 3 modules captured different tumor sub-regions and one module was mainly expressed in the stromal connective tissue (Fig. 3A). The combination of the relative intensities of the MEs localized within the tumor region revealed the complex spatial patterns associated with its molecular heterogeneity (Fig 3A). The module preservation analysis (5,000 permutations) showed that 4 of the 6 metastatic network modules were from weakly to moderately preserved in the *X_non-met_* network. Among those, the green module was associated with tissue and was weakly preserved (*Z_summary_* ≾ 5) in the *X_non-met_* data (Fig. S1A). The relatively lower correlation (Pearson’s r = 0.65) between the 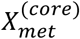 and 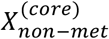 module membership (kME), defined as the correlation between the ions intensities and the ME, confirmed that the green module ions were not equally co-localized in the two datasets (Fig. S1A). The projection of its ME intensities on the optical image of the H…E stained tissue showed that the green module ions involved the stromal tissue segment of the tumor with the observed presence of free fat droplets and infiltrations of tumor cells (an example is reported in Fig 4A). Noticeably, the MEs revealed that the green module ions were also expressed in the 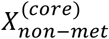 samples (Fig. S3) confirming that the relevant biochemical differences could be captured only in terms of different molecular interactions (network edge weights) and not in terms of presence/absence of certain sets of ions. Analyzing the corresponding regions in the optical images of the H…E stained tissue, it was observed that this module corresponded to regions close to the necrotic tissue (an example is reported in Fig. 4B).

**Fig. 2.**
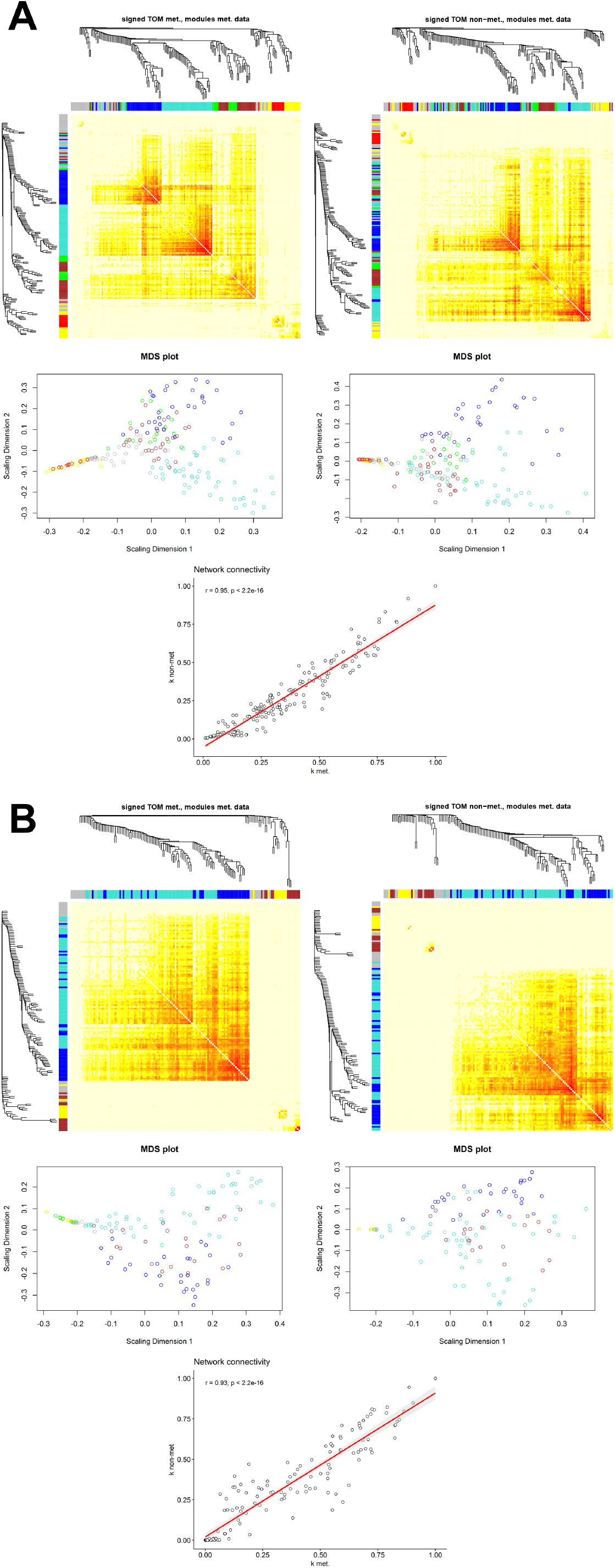
*– Graphical representation of the co-expression networks. Panel A: TOM heatmap with hierarchical clustering results for the metastatic (top left) and non-metastatic (top right) tumor center data; multidimensional scaling (MDS) scatter plot showing the node distribution associated with the module color attributes for the metastatic (center left) and non-metastatic (center right) tumor center data. The high Pearson’s correlation value between the network connectivity k of the metastatic and non-metastatic tumor center data confirms that the two networks are characterized by similar global topological properties. Panel B: Analogous results for the data associated with the tissue sections at a distance of 10cm from the tumor. Figures at left represent the results from the metastatic samples, and figures at right represent the results from non-metastatic samples*.

**Fig. 3.**
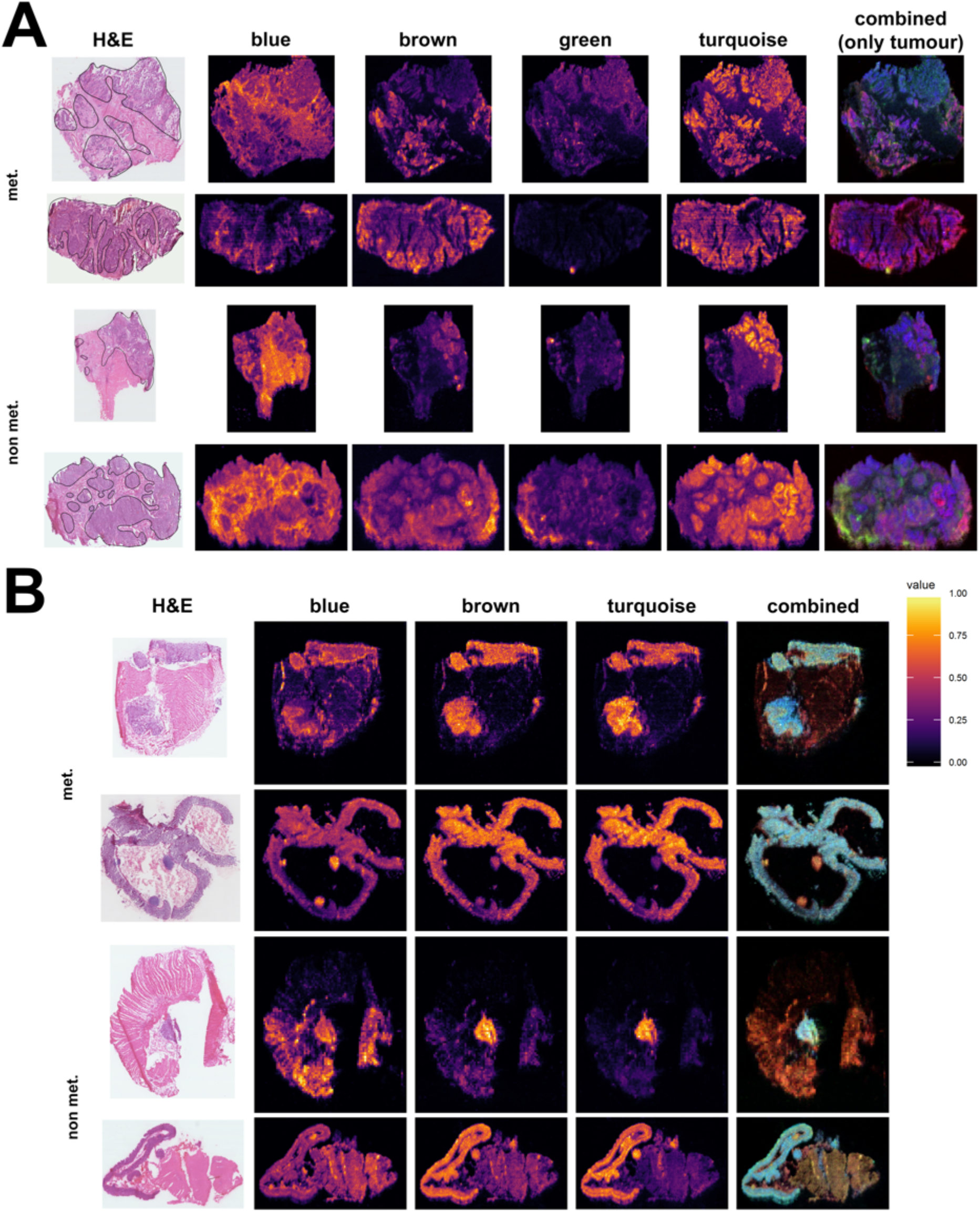
*– Spatial visualization of the tissue related module eigenmetabolites (ME). Panel A: results for the tumor core samples. The four tissue related modules are localized in different regions of the cancerous tissue, the “blue” module is mainly expressed in the connective tissue surrounding the tumor (delineated by a black line in the H…E image in the first column), whereas the other three modules mostly involve the tumor. The tumor molecular heterogeneity is captured by the modules “brown”, “green” and “turquoise” as shown by the combination of their ME in the last column. Here the three ME expressions are scaled in [0, 1] and visualized as the intensities of RGB channels. Two examples are reported for both the metastatic and non-metastatic samples. Panel B: Analogous results for the tissue i section at 10cm from the tumor center. Here the three tissue related modules are mainly expressed in) the epithelial tissue, as shown by the H…E in the first column. The analogous combination of the ME*) *into a RGB image shows the molecular heterogeneity captured by WGCNA*.

**Fig. 4.**
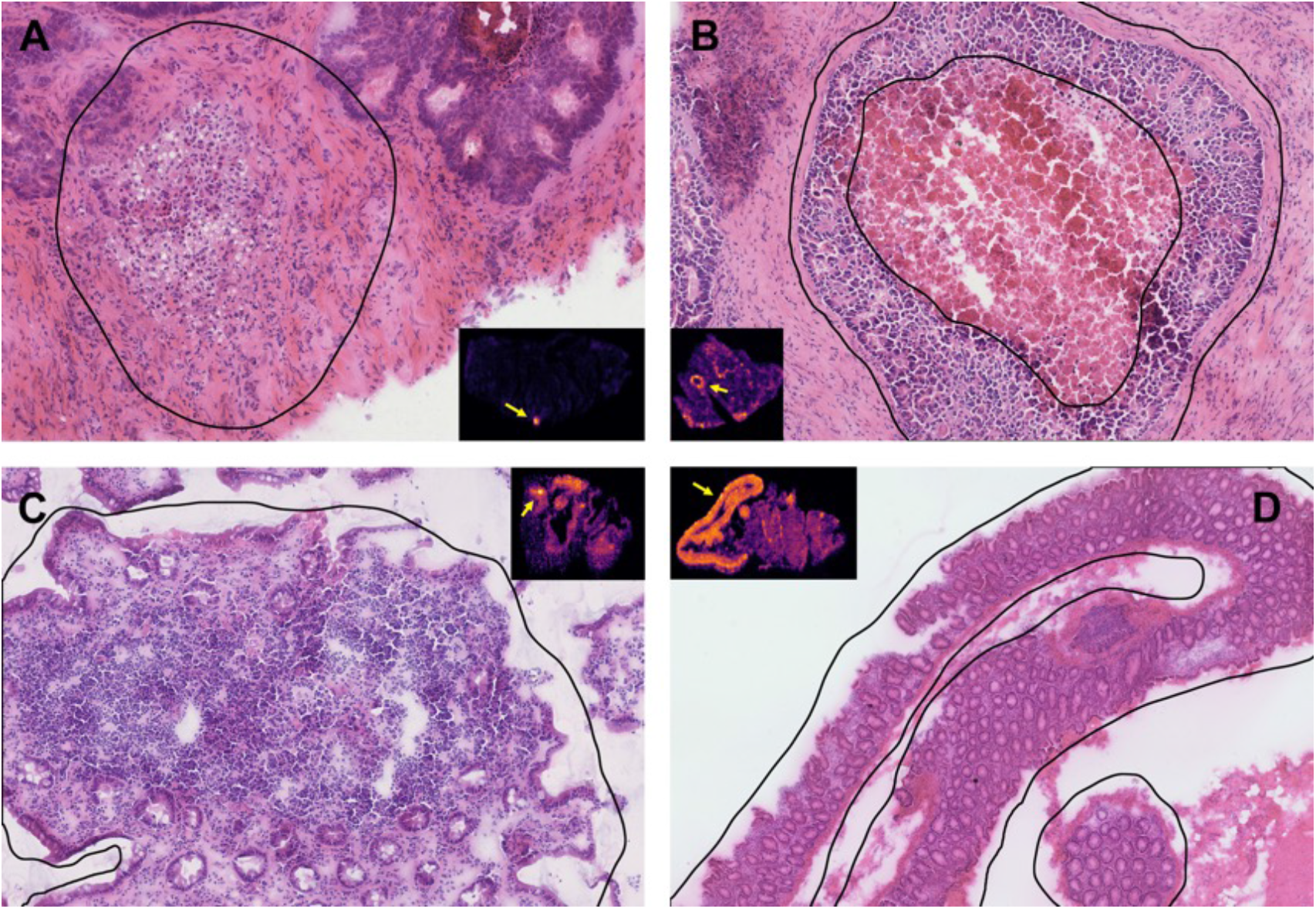
*– Examples of the observed tissue associated with the identified modules. In the metastatic tumor center (A), the weakly preserved “green” module ions are localized in the stroma surrounding the tumor. Noticeably macrophages infiltrating the area are visible together with free fat droplets. The latter have been previously associated with an inflammatory response to the infiltration of the tumor cells (Bozza and Viola 2010). In the non-metastatic tumor core, the module ions are still associated with an inflammatory condition that seems to be driven by a necrosis process, as shown in the zoomed H…E stained image corresponding region of the module eigenmetabolites image pointed by the arrow (small box) (B). This observations suggest that different mechanisms are driving the local inflammatory response. Analogously, at a distance of 10cm from the tumor, the weakly preserved “brown” module is localized in sub regions of the metastatic related epithelium involving groups of lymphatic cells as shown by the highly dense cellular populations in the zoomed H…E stained image corresponding to the region pointed by the arrow in the module eigenmetabolites image (small box) (C). A similar result can be observed in the non-metastatic related tissue (D), as represented by the zoomed region of H…E corresponding to the higher intense module eigenmetabolites region. Here, clusters of lymphatic cells are involved in the selected regions of the non-metastatic tissue as well*.

The ion annotation, performed through a Metlin database search (Smith, O’Maille et al. 2005), revealed that those of the weakly preserved green module consisted of phosphatidylglycerols (PG) and fatty acids (FA) (Fig 5A, Table S1). Specifically, hexadecanoic acid (C16:0), eicosatetraenoic acid (C20:4), docosatetraenoic acid (C22:4), docosapentaenoic acid (C22:5), docosahexaenoic acid (C22:6) and tetracosapentaenoic acid (C24:5) which showed high correlation with the distribution of PG(36:3), PG(36:4), PG(38:5), PG(38:6), PG(40:6), PG(40:7) and the non-annotated 292.2432, 528.2736 and 791.5441 *m/z* ions. The ion pair specificity (SI Appendix, Materials and Methods), confirmed that most of the PGs together with tetracosapentaenoic acid were specific of the 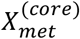 sample set (Fig. 5A).

**Fig. 5.**
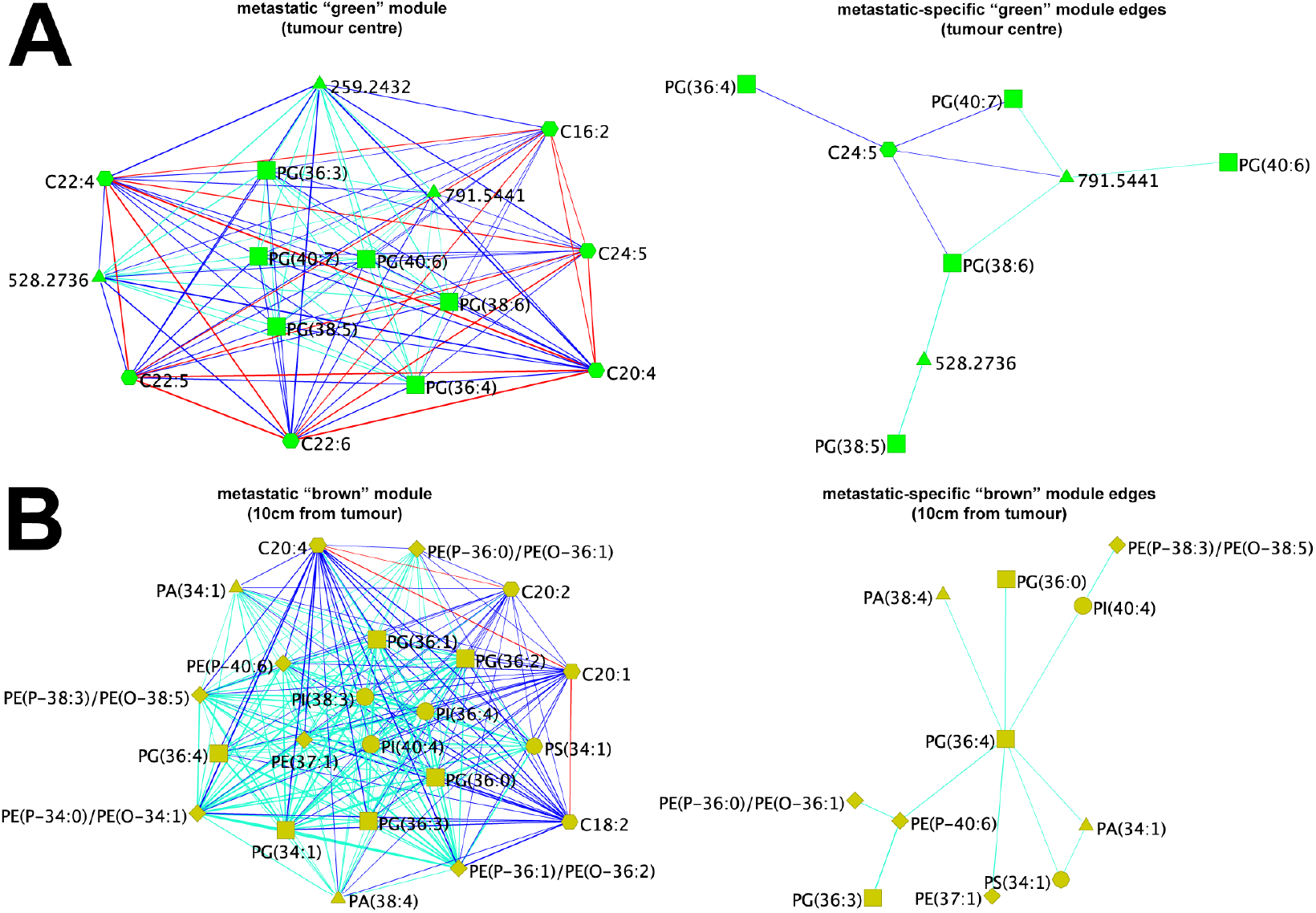
*– Graphical representation of the metastatic related modules that are weakly preserved in the non-metastatic networks. The “green” module associated with the tumor core tissue (panel A, left) consists of PG and PUFA. The edges connecting the PUFA and the PG are colored in blue, whereas those connecting PUFA to PUFA are colored in red. A metastatic edge specificity larger than 0.8 reveals that PG and C24:5 edges are present in the metastatic network but absent in the non-metastatic network (A, right). Moreover, the higher number of edges involving C24:5 and the ion 791.5441 m/z makes them the most representative ions of the metastatic-related module. Analogously, the weakly preserved “brown” module of the network associated with the metastatic 10cm tissue sections reveals a more complex co-localization pattern involving different families of phospholipids and fatty acids (panel B, left). In particular, the metastatic-specific module edges ions, in this case, involve mainly phospholipids, with a central role played by PG(36:4) (panel B, right)*.

Confirmation of the different network properties between the discovery and test datasets was obtained by running WGCNA directly on the 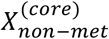 samples. Indeed, it was seen that the green module ions were split into two different non-metastatic modules (Fig. S4C). Most of the FAs were assigned to a different module together with phosphatidylethanolamines (PEs), phosphatidylinositols (PIs) and phosphoserines (PSs) (Table S2, Fig. S4D). The PGs were instead assigned together in a larger module consisting of only phospholipids (Table S3, Fig. S4E).

### Phospholipid metabolism is different at 10cm from the tumor core of metastatic and non-metastatic patients

Analogous to the tumor core analysis, module preservation analysis was performed using the MSI data from tissue samples collected from surgical specimens at 10cm from the tumor core. The specimens from 29 patients (Table 1) were split into two groups corresponding to the occurrence/nonoccurrence of a metastatic relapse during the follow-up period. The MSI data of the 6 patients who developed metastases, denoted 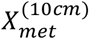, were used to determine the reference network modules that were later tested for preservation in the remaining 23 patients MSI data 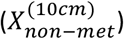. As seen in the tumor center samples, the WGCNA consensus networks calculated from the mass spectral data from 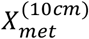 and 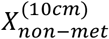 were characterized by similar global properties (Pearson’s r = 0.93 between metastatic and non-metastatic data k, Fig. 2B); this result also showed that in this case the two metabolic networks were globally almost identical. The metastatic related network was clustered in 6 modules (denoted “blue”, “brown”, “green”, “turquoise”, and “yellow”) (Fig 2B). Analyzing the spatial distribution of the detected ME, it was observed that three of those modules showed were localized in the tissue of all 6 patients (Fig. S5). The combination of the relative intensity of those MEs revealed the molecular heterogeneity of the tissue sections (Fig. 3B). Using module preservation analysis with 5,000 permutations, it was observed that all the modules were from weakly to moderately preserved (2 ≤ *Z_summary_* ≤ 10). In particular, the brown module, expressed in the tissue, was weakly preserved (*Z_summary_* ≾ 5) in the patients who did not develop metastasis. As observed in the tumor core data, the 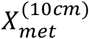 and 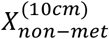 the brown module *kMEs* showed a relatively lower correlation (Pearson’s r = 0.74, Fig. S1B).

The annotation of the brown module ions, mainly expressed in the epithelium (as revealed by a comparison with the optical images of the H…E stained images, shown in Fig. 4C-D), consisted of different classes of phospholipids (PE, PG, PI, and PS) together with phosphatidic acids (PA), octadecadienoic acid (C18:2), eicosenoic acid (C20:1), eicosadienoic acid (C20:2), and eicosatetraenoic acid (C20:4) (Table S4). The ion pair significance showed that the phospholipid distributions showed stronger correlations in 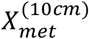 than in 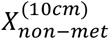 samples (Fig 5B), with PG(36:4) playing a central role in the metastatic network specific edges.

Running WGCNA directly on the 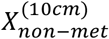 MSI data showed that most of the brown module ions were assigned to a more complex module, labelled “turquoise”, together with different classes of phospholipids and FAs (Fig. S7A-C). Specifically, among the annotated ions, the most probable candidates for 281.2486, 279.233, 277.2174, 309.28, 306.2521 *m/z* were identified to be n-9 oleic acid (C18:1), n-6 linoleic acid (C18:2), n-3/6 α/γ-linoleic acid (C18:3), n-9 eicosenoic acid (C20:1), n-6 dihomo-γ-linoleic acid (C20:3) and n-3 docosahexaenoic acid (C22:6); those ions were found colocalised with the phospholipids (Table S5, Fig. S7D).

## Discussion

Understanding the mechanisms behind the heterogeneity of cancerous tissues is a key goal for the development of effective cancer treatment strategies (Murugaesu, Chew et al. 2013). In this study, a network based analysis of the co-localized ions allowed the identification of a set of tissue sub-types (ion modules) that represent the tissue heterogeneity associated with each individual subject. The WGCNA approach, combined with the Dynamic Tree Cut algorithm, determined the optimal number of tissue sub-types with no user intervention. The only parameter required, the minimum number of ions necessary to form a module, was set to 5 following the hypothesis that small clusters could capture finer details of the heterogeneous metabolic patterns. In order to detect the metabolic differences between the group-of-interest (patients developing metastasis) and the remaining part of the cohort, the modules detected were tested in the former for preservation in the latter group. The permutation test applied on MSI data at a distance of 10cm from the tumor core allowed the identification of one module associated with the development of metastasis in both the cases.

A set of molecular interactions involving PG and FAs was found to be representative of the increased risk of the development of metastases in the tumor core. Specifically, the combination of WGCNA and the differential network analysis showed that, in the metastatic tumor core set, PGs were more highly co-localized together with tetracosapentaenoic acid than in the non-metastatic related samples, suggesting a key role for these molecules in differentiating the local biochemical reactions. The presence of polyunsaturated fatty acids (PUFAs) in the “brown” module together with PGs suggests that these regions elicit significantly higher phospholipase A2 (PLA2) activity. This enables the formation of prostaglandins, in particular prostaglandin E2, through COX2 activity, which has widely been associated with tumor invasion and metastasis formation in a broad range of different cancers. cPLA2 activity itself has been found to contribute to metastasis formation in breast cancer *via* TGF β- induced epithelial-mesenchymal transition (EMT) (Chen, Fu et al. 2017).

PGs are generally considered to be membrane constituents of bacterial cells, serving mainly as surfactants in mammalian organisms. In this regard, the PGs detected are unlikely to be building blocks of cellular membranes in the tumor environment, but rather present in the interstitial space. Considering the apparent specificity of PLA2 for anionic phospholipids likely excreted into the extracellular space and the rich abundance of macrophages in association with the green module (Fig. 4A), the underlying enzyme is probably the PLA2 produced by macrophages. Macrophage cPLA2 has already been associated with metastasis formation in case of lung cancer and it was successfully demonstrated that depletion of macrophage cPLA2 by gene knock out resulted in the significant decrease of metastatic potential (Weiser-Evans, Wang et al. 2009). In conclusion, the macrophage infiltration clearly shown on Fig. 4A is likely to confer sufficient PLA2 activity to trigger EMT and downstream metastasis formation in the corresponding group. Although the involvement of gut microbiota is not obvious in the current case, bacterial PGs, as the main constituents of prokaryotic cellular membranes can potentially contribute to the expression of macrophageal PLA2 through TLR2 pathway, especially in the proximity of necrotic areas.

From analysis of the weakly preserved module between the metastatic and non-metastatic related tissue samples at 10cm from the tumor core, it was found that the PUFAs octadecadienoic acid, eicosenoic acid, eicosadienoic acid and eicosatetraenoic acid were correlated mainly with PG, PE and plasmalogen PE together with PS and PI in sub-regions of epithelial tissue. In this case, the significantly stronger co-expression levels of the phospholipids determined by their high significance in the metastatic related tissues can be interpreted as the result of increased activity of a number of enzymes including phospholipase D (PLD). Abnormalities in PLD expression can be responsible for altered cell proliferation mechanisms (Foster and Xu 2003, Park, Lee et al. 2012). Similarly to the tumor core, a relatively high correlation of eicosatetraenoic acid with most of the detected phospholipids was observed in the weakly preserved module, suggesting evidence of PLA2 activity that can be interpreted as a localized inflammatory condition, also confirmed by the visual inspection of the corresponding regions in the H…E stained tissues (Fig 4B). This, combined with the abundance of plasmalogens in the module, implies peroxisomal involvement in metastasis formation. It has recently been demonstrated that peroxisomal functionality is key for macrophage activation (Di Cara, Sheshachalam et al. 2017) which can explain the correlation between the peroxisomal metabolic phenotype represented by the network structures (Fig 5B) and metastatic potential conferred by macrophage PLA2 activity.

## Conclusions

Tumors are characterized by a wide range of metabolic, genetic, and phenotypic properties. This heterogeneity, expressed at the genetic and epigenetic level, is reflected by the metabolites produced within cells. In this study, DESI-MSI was shown to be effective for untargeted analysis of the local metabolism of colorectal cancer tissue. The differences in the correlations between the distribution of key ions and their preservation in the network modules identified that the increased activity of PLA2, exploiting the high availability of PG in the surfactant layer in the tumor center, was associated with a risk of post-surgical distant metastasis formation. The molecular correlations which were not preserved revealed that eicosanoid precursors in stromal tissue can play a significant role in the increased cellular proliferation within the tumor core. At 10cm from the tumor center, the altered peroxisomal metabolic phenotype was found to be associated with the different metastatic potential, likely *via* a macrophage activation mechanism. Finally, the network approach identified a complex picture of tumor heterogeneity, confirming that a simple ‘presence/absence of ions’ approach is not able to determine the differences between the metabolic pathways.

Due to the limited number of patients considered, these results should be considered preliminary and await confirmation in a larger patient cohort. In future work, the study will be expanded to a larger number of patients, allowing a more detailed analysis of the metabolic differences associated with a range of clinical characteristics. Furthermore, MSI data in combination with local genetic expression identified from spatial ME maps will be included in order to investigate both genetic and metabolic changes to achieve deeper insight of the nature of tumor heterogeneity and its chemical/metabolic interactions with the surrounding tissue.

## Materials and Methods

DESI imaging data was acquired from sections of individual tumor central regions and from section of tissue at 10cm from the tumor central regions of, respectively, 32 and 29 subjects affected by colorectal cancer (SI Appendix, Materials and Methods). The tissue specimens were collected after the surgical removal of the tumor. The subjects were monitored after surgery for a period of up to 6 years, and events such as disease occurrences (metastases, or local recurrence) and deaths were recorded. During the observational period, 8 of the 32 subjects and 6 of the 29 subjects developed metastatic recurrence. The tissue sections were fresh-frozen and mass spectrometry imaging (MSI) data were acquired using an automated 2D DESI source mounted to a Thermo Exactive Orbitrap mass spectrometer. MS data were acquired in the negative ion mode, in the range of 200-1,050 *m/z*. The RAW spectra, corresponding to tissue regions (pixels) were pre-processed in order to extract a set of variables associated with the informative peaks and their relative abundances. Centroided data were extracted using the ProteoWizard software (Kessner, Chambers et al. 2008) and pre-processed using the ‘MALDIquant’ package for R (Ihaka and Gentleman 1996, Gibb and Strimmer 2012). Before preprocessing, a re-calibration procedure was applied in order to reduce the peak shift across the pixels of each tissue sample. The procedure is described in the ‘Results’ section. The details of the preprocessing workflow are reported in SI Appendix, Materials and Methods. The tumor core preprocessed data corresponded to a series of matrices (pixels x ions) containing the intensities of the 185 *m/z* values common to all the sample pixels. For the samples at 10cm from the tumor, 141 *m/z* values were found common to all the samples. The consensus adjacency matrix elements were defined as the minimum of the ions pairwise Pearson’s correlations associated with each tissue section. In a similar fashion, the TOM was calculated using the signed adjacency raised to the soft power value. Modules were identified by applying the Dynamic Tree Cut hybrid algorithm to the hierarchical dendrogram with an average linkage obtained using *1-TOM* as distance matrix. Module preservation was performed using the WGCNA *‘modulePreservatiorí* command with 5000 permutations, using the consensus adjacency matrix as input. The module identification procedure was applied directly on the consensus matrices of the non-metastatic related MSI data for both the tumor core and 10cm datasets.

Datasets and scripts are publicly available at https://doi.org/10.4121/uuid:f06dee0d-1d2e-4d67-b978-5bac4087d346.

## Acknowledgements

Full ethical approval was obtained from the institutional review board at Imperial College Healthcare NHS Trust (REC reference numbers 07/H0712/112, 11/LO/0686). The project was funded by the European Research Council (ERC Consolidator Grant ‘MASSLIP’ Contract No: 617896), the National Institute for Health Research through the Imperial Biomedical Research Centre and Cancer Research UK Grand Challenge Program (Project title: “A Complete Cartography Through Multiscale Molecular Imaging”).

